# Single cell transcriptomic re-analysis of immune cells in bronchoalveolar lavage fluids reveals the correlation of B cell characteristics and disease severity of patients with SARS-CoV-2 infection

**DOI:** 10.1101/2020.11.09.374272

**Authors:** Chae Won Kim, Ji Eun Oh, Heung Kyu Lee

## Abstract

The COVID-19 pandemic (SARS-CoV-2) is a global infectious disease with rapid spread. Some patients have severe symptoms and clinical signs caused by an excessive inflammatory response, which increases the risk of mortality. In this study, we reanalyzed scRNA-seq data of cells from bronchoalveolar lavage fluids of patients with COVID-19 with mild and severe symptoms, focusing on antibody-producing cells. In patients with severe disease, B cells seemed to be more activated and expressed more immunoglobulin genes compared with cells from patients with mild disease, and macrophages expressed higher levels of the TNF superfamily member B-cell activating factor but not of APRIL (a proliferation-inducing ligand). In addition, macrophages from patients with severe disease had increased pro-inflammatory features and pathways associated with Fc receptor-mediated signaling, compared with patients with mild disease. CCR2-positive plasma cells accumulated in patients with severe disease, probably because of increased CCL2 expression on macrophages from patients with severe disease. Together, these results support that different characteristics of B cells might affect the severity of COVID-19 infection.

## Introduction

The COVID-19 pandemic is a global infectious disease caused by severe acute respiratory syndrome coronavirus 2 (SARS-CoV-2). The first reports of the virus and associated disease were cases in Wuhan, China in December 2019 (1). As of June 22, 2020, there were 8,860,331 individuals worldwide with confirmed infections, with a mortality rate of approximately 5% (2). Some patients with severe symptoms and clinical signs experience acute respiratory distress syndrome or hyper-inflammatory syndrome, which increases mortality (3). To reduce COVID-19-associated mortality rates, it is necessary to understand the mechanisms of hyper-inflammatory syndrome and develop efficient treatments that block the effects.

To characterize the immunologic signatures differed with severity of COVID-19 disease, recent studies showed transcriptomic analysis of immune cells, especially myeloid cells, from SARS-CoV-2-infected patients. In patients with severe COVID-19, not with milder COVID-19, classical monocytes showed type I interferon (IFN) response together with tumor necrosis factor/interleukin-1β (TNF/IL-1β)-driven inflammation (4). Another transcriptomic study with samples from COVID-19 patients showed that monocyte-derived macrophages in bronchoalveolar lavage (BAL) fluid from patients with severe COVID-19 have pro-inflammatory features (5). In addition, the other study observed reduced expression of human leukocyte antigen class DR (HLA-DR) and type I IFN deficiency in the myeloid cells of patients with severe COVID-19 (6). Furthermore, in patients with severe COVID-19, inflammatory macrophages expressing pro-inflammatory cytokines and chemokines were observed (7).

B cells, as well as myeloid cell, play important roles in antiviral immune responses via inducing humoral responses. Indeed, neutralizing antibodies against SARS-CoV-2 are able to regulate infection in vitro and in vivo (8–10). While titers of IgG specific to spike protein were maintained in recovered patients, not all of them have detectable neutralizing antibodies (11). Furthermore, next-generation sequencing of B cell receptor (BCR) repertoires from COVID-19 patients showed the association between somatic hypermutation and a more severe clinical state (12). Some studies have revealed associations between higher titers of antibodies and severe COVID-19 (1,13), suggesting that antibody-dependent enhancement (ADE) is one of the possible mechanisms triggering severe macrophage-associated inflammation during SARS-CoV-2 infection. Therefore, understanding B cells in SARS-CoV-2 infection situation is one of critical points for developing COVID-19 therapies. Despite the importance of humoral responses, the characteristic of B cells in lung with the severity of COVID-19 patients was still unclear. In this study, to find out the features of B cells that differ with severity of COVID-19, we analyzed data from single-cell RNA sequencing (scRNA-seq) results for cells from BAL fluids of patients with mild and severe COVID-19.

## Materials and Methods

### Single-Cell RNA Sequencing

A publicly available data set was used for the analysis (Gene Expression Omnibus database, GSE145926). We used scRNA-seq data from BAL fluids of healthy people (control group) and from patients with COVID-19 with mild and severe symptoms and clinical signs (healthy control = 3, mild = 3, severe = 6) (Table S1). The analysis was performed using the Seurat R software package (version 3.1.5). We analyzed cells for which the features were between 200 and 7500, and mitochondrial gene percentages were < 10–25%. The filtered cells were normalized using the ‘LogNormalize’ method (scale.factor = 10000). The top 2000 of the variable genes were identified using the ‘vst’ method in the FindVariableFeatures function. All samples were then integrated into a matrix using FindIntegrationAnchors and IntegrateData functions with 50 dimensions. The matrix was scaled using the ScaleData function (default setting), and principal-component analysis was performed using 50 principal components. Finally, uniform manifold approximation and projection was performed based on 50 dimensions. The cells were clustered using the FindClusters function (resolution = 1.5).

### Visualization of Gene Expression

The idents of the clusters were renamed into cell type, and clusters with the same idents were combined. The B cells were re-clustered based on seven dimensions (resolution = 0.5). Violin plots and feature plots were used to visualize expression of the genes of interest for each cell type. Dot plots were also used to visualize expression of several genes.

### Differential Gene Expression Analysis

To compare the differential gene expression in macrophages, B cells, and plasma cells between patients with mild and severe disease, the FindMarkers function was used identify differential expression genes (DEGs). In this function, upregulated genes in patients with mild disease compared with patients with severe disease were specified as positive or negative, respectively. The list of DEGs was ranked by adjusted p-value.

### Gene Set Enrichment Analysis

Gene set enrichment analysis (GSEA) software (GSEA, version 4.0.3) was used for the analysis. The lists of DEGs were exported into a ranked file format and were uploaded to the GSEA software. The analysis was based on 1000 permutations; Symbol with Remapping in the Molecular Signatures Database reference platform (version 7.1) was used for DEG annotation. HALLMARK, the Kyoto Encyclopedia of Genes and Genomes (KEGG), and Gene Ontology (GO) analysis were used for analysis of macrophages. The plasma cell data were analyzed using GO analysis. The B cell data were analyzed using HALLMARK, GO, and immunological signature analyses.

## Results

### B Cells Appear to be More Activated in Patients with Severe Disease

We analyzed scRNA-seq data of cells from BAL fluids of patients with mild (n=3) or severe (n=6) COVID-19 and of healthy controls (n=3) (5) because lung inflammation could be critical for disease pathogenesis. Using cluster analysis, we acquired 34 clusters of BAL fluid cells (Figure S1A) and annotated them according to the gene markers (5); Macrophage (*CD68, HLA-DRA*), T cell (*CD3D*), Neutrophil (*FCGR3B*), Epithelial cell (*TPPP3, KRT18*), NK cell (*KLRD1*), Dendritic cell (*HLA-DRA, CD1C, CLEC9A*), Plasma cell (*IGHG4*), B cell (*MS4A1*), Plasmacytoid dendritic cell (*LILRA4*), and Mast cell (*TPSB2*) (Figure S1B). Cluster 27 was excluded because both *CD3D* and *CD68* were expressed. The distributions of all cell types were different for each group (Figure S1C).

At viral infection, B cells are activated and differentiated into plasma cells producing antibodies against viral antigens. We analyzed B cells in the BAL fluids from the three groups (mild, severe, control) to determine the characteristics of B cells. The results indicated that the proportion of B cells was increased in mild disease group, compared with the healthy group. While the proportion of B cells was lower in the severe group compared with the mild group, the actual count of B cells was elevated in the severe group (Figure 1A). To compare characteristics of the B cells between the mild and severe groups, we performed GSEA for B cells using gene sets related to B cell differentiation. The results indicated that the upregulated genes in the B cells from the patients with mild disease were relatively more associated with naïve B cells or memory B cells, rather than plasma cells (Figure 1B). Because B cells express different chemokine receptors depending on different subsets and activation status (14,15), we analyzed expression of chemokine receptor genes on B cells. B cells from the patients with mild disease mainly expressed *CCR6* and *CXCR3*, but those from patients with severe disease highly expressed *CXCR4* (Figure 1C). These results indicated that the characteristics of the B cells between the two groups were different, and that the characteristics of the B cells from patients with mild disease were more associated with memory B cells.

**Figure 1.**
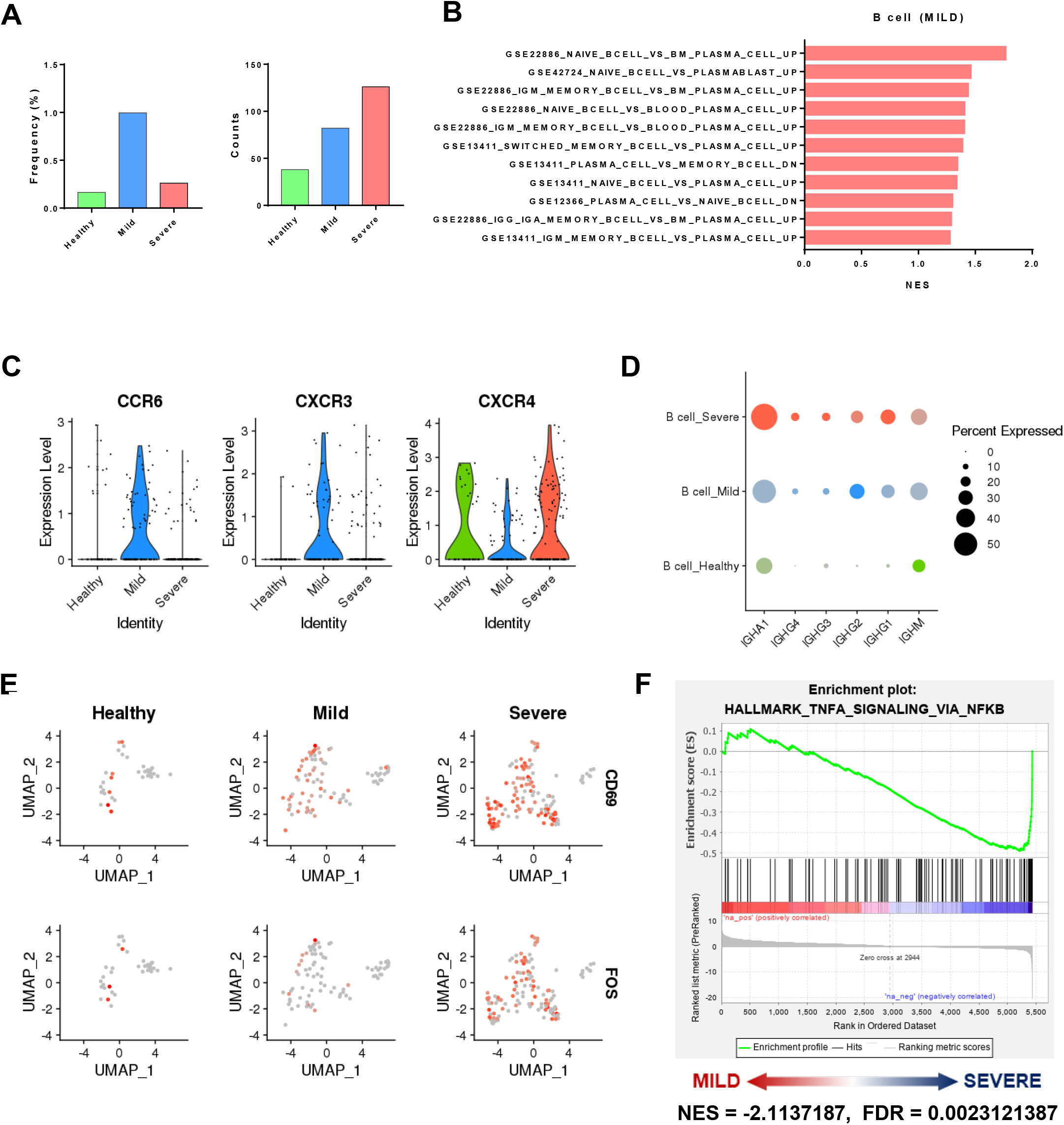
Different characteristics of B cells in patients with mild or severe COVID-19. (A) Proportions (left) and counts (right) of B cells among healthy controls and patients with mild or severe SARS-CoV-2 infection. (B) Immunologic signature analysis related to B cells and plasma cells of DEGs in B cells of patients with severe, compared those from patients with mild, disease, top 10 highlighted gene sets. (C) Expression of *CCR6* and *CXCR4* on B cells from each sample. (D) Expression of immunoglobulin genes (*IGHM, IGHG1, IGHG2, IGHG3, IGHG4, IGHA1*) on B cells from each sample (E) UMAP plots of *CD69* and *FOS* gene expression on B cells of healthy controls and patients with mild or severe infection. (F) GSEA plot of DEGs in B cells of patients with mild vs severe COVID-19, HALLMARK_TNFA_SIGNALING_VIA_NFKB gene set

After activation, immunoglobulin (Ig) class switching occurs that results in changes in expression of IgM and IgD into IgG, IgA, or IgE *(16)*. Expression of immunoglobulin genes such as IGHG1, IGHG2, IGHG3, IGHG4, and IGHA1 were increased in B cells from the patients with severe disease compared with the patients with mild disease (Figure 1D). These results suggested that the B cells from the severe group were activated and experienced an antibody-switching process. Moreover, expression of CD69 and FOS in B cells, which are upregulated upon activaiton *(17,18)*, was higher in the severe group than in the mild group (Figure 1E). To determine which pathways affected activation of B cells from the patients with severe disease, we performed GSEA with HALLMARK gene sets and acquired significant results for tumor necrosis factor (TNF)-α signaling (Figure 1F). These results suggested that the B cells from the patients with severe disease were more activated by TNF-α signaling, compared with those from the patients with mild disease.

### Expression of APRIL, not B-Cell Activating Factor, on Macrophages is Decreased in Patients with Severe COVID-19

In addition to TNF-α, B-cell activating factor (BAFF) and a proliferation-inducing ligand (APRIL) are members of the TNF superfamily, which are involved in activation and differentiation of B cells (19). Since the GSEA result based on the TNF-α signaling gene set indicated that upregulated genes of B cells in the severe group were more associated with TNF-α signaling than in the mild group (Figure 1F), it is possible that BAFF and APRIL also affected B cells during SARS-CoV-2 infection.

There are three known receptors for BAFF and APRIL (i.e., BAFF receptor (BAFFR, encoded by *TNFRSF13C*), transmembrane activator and calcium modulator and cyclophilin ligand interactor (TACI, encoded by *TNFRSF13B*), and B cell maturation antigen (BCMA, encoded by *TNFRSF17*)). BAFF binds to all three receptors, but APRIL binds to TACI and BCMA (20). The results of the analysis indicated that B cells from patients with mild or severe disease expressed both genes encoding BAFFR and TACI, but gene encoding BCMA was only slightly expressed (Figure S2A). These results suggested that BAFF and APRIL can affect B cells through those receptors.

We analyzed expression of *TNFSF13B* (encoding BAFF) and *TNFSF13* (encoding APRIL) on the cells from the BAL fluids of the patients with COVID-19 (Figure 2A). *TNFSF13B* was highly expressed by macrophages and neutrophils, whereas *TNFSF13* was expressed only by macrophages. We then compared expression of two genes in the neutrophils and macrophages among the three groups. Although accumulation of neutrophils occurred in BAL fluids from the patients with severe disease (Figure S1D), there were no differences in *TNFSF13B* and *TNFSF13* expression in neutrophils between the mild and severe group (Figure S2B). Expression of *TNFSF13B* in macrophages seemed slightly higher in the severe group, while expression of *TNFSF13* was much lower in the severe group compared to the mild group (Figure 2B, S2C).

**Figure 2.**
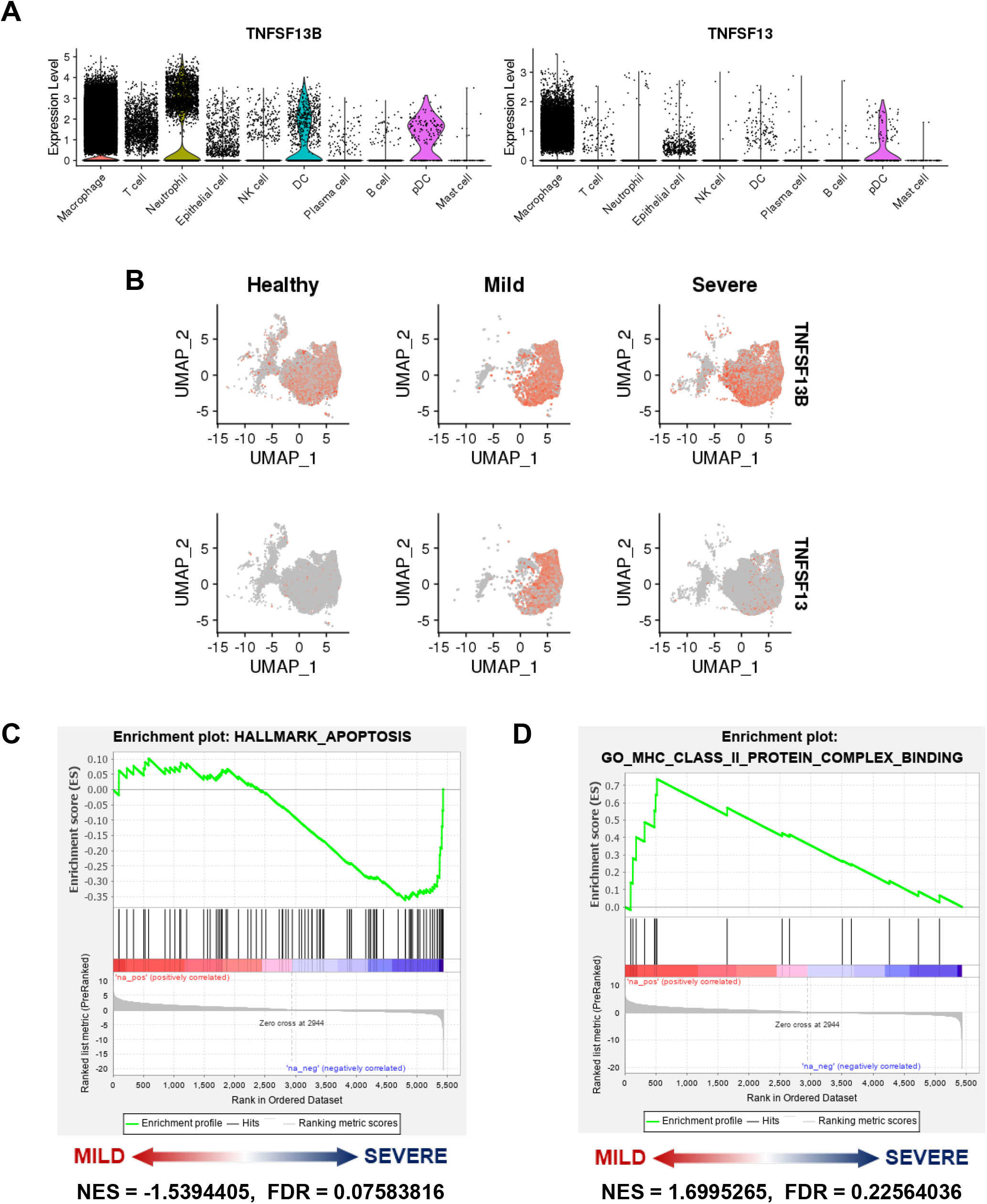
Decreased APRIL expression of macrophages in patients with severe COVID-19. (A) Expression of *BAFF* (*TNFSF13B*) and *APRIL* (*TNFSF13*) on each cell type. (B) UMAP plots of *TNFSF13B* and *TNFSF13* on macrophages from healthy controls and from patients with mild or severe disease. (C, D) GSEA plot of DEGs in B cells of patients with mild vs severe disease, with (C) the HALLMARK_APOPTOSIS gene set and (D) the GO_MHC_CLASS_II_PROTEIN_ COMPLEX_BINDING gene set.

To examine the effects of BAFF and APRIL on B cells, we performed GSEA of B cells from BAL fluids of the patients with mild and severe disease. While BAFF can promote B cell survival and growth through BAFF-R signaling, it can induce B cell apoptosis via TACI signaling (21). The GSEA result indicated that the B cells from severe group had more genes upregulated that were related to apoptosis (Figure 2C). APRIL promotes antigen presentation of B cells through BCMA signaling (22). The GSEA also revealed that upregulation of genes related to major histocompatibility complex class II protein complex binding occurred more in the B cells from patients with severe disease (Figure 2D). Taken together, these results suggested that the B cells from the patients in the severe group were more affected by BAFF and less by APRIL.

### Macrophages from Patients with Severe COVID-19 Have More Pro-Inflammatory Features, Especially Mediated by Antibody-dependent Signaling

The analysis revealed that macrophages were the most abundant of all the cell types in the BAL fluids (Figure S1D). To confirm the pro-inflammatory features of macrophages associated with SARS-CoV-2 infection reported in previous study (5), we analyzed expression of *IL6* and *TNF*, which trigger cytokine storms during COVID-19 (23). The results for the feature plots of the two cytokine genes indicated that compared with the healthy controls, expression was induced in patients with the infection and was higher in patients with severe symptoms and clinical signs (Figure 3A). In addition to *IL6* and *TNF*, expression of *IL1B* and *CCL2* (i.e., pro-inflammatory indicators) were much higher on macrophages isolated from patients with severe disease than from patients with mild disease (Figure 3B). The GSEA revealed that the upregulated genes expressed on macrophages from patients with severe disease were more related to the inflammatory response than those from patients with mild COVID-19 (Figure 1C).

**Figure 3.**
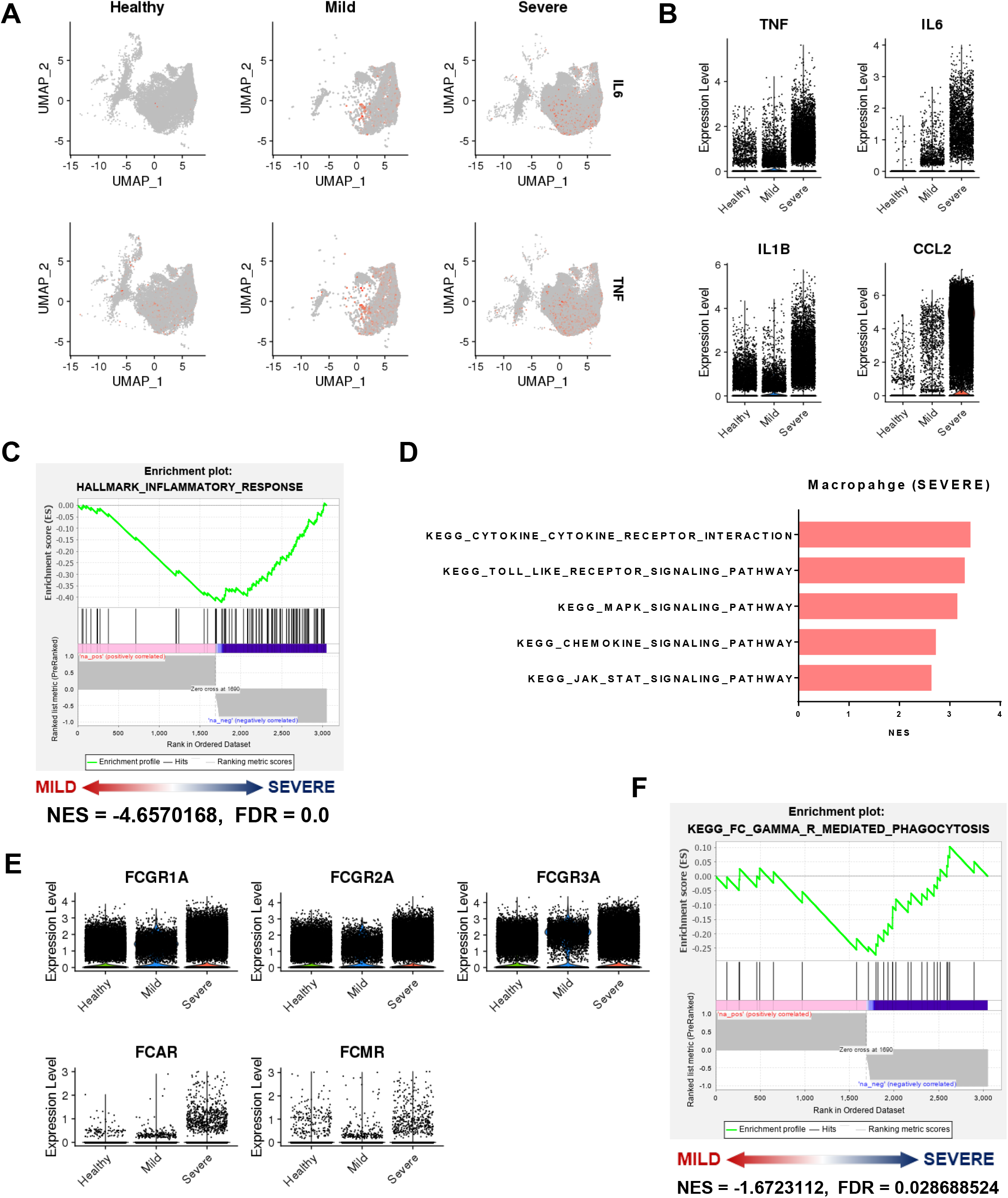
Macrophages from patients with COVID-19 have pro-inflammatory features. (A) Uniform manifold approximation and projection (UMAP) plots showing *IL6* and *TNF* gene expression on macrophages from bronchoalveolar lavage fluids from healthy controls and from patients with mild or severe COVID-19. (B) Expression of pro-inflammatory cytokines and chemokines (*TNF, IL6, IL1B,* and *CCL2*) on macrophages from each sample. (C) GSEA plot of DEGs in macrophages from patients with mild vs severe disease, HALLMARK_ INFLAMMATORY_RESPONSE gene set. (D) KEGG analysis of up-regulated DEGs in macrophages from patients with severe, compared with those with mild, disease showing the top five highlighted pathways. (E) GSEA plot of DEGs in macrophages from patients with mild vs severe COVID-19, KEGG_FC_GAMMA_R_MEDIATED_PHAGOCYTOSIS gene set. (F) Expression of activating Fc gamma receptor gene (*FCGR1A, FCGR2A, FCGR3A, FCAR, FCMR*) on macrophages from each sample.

Macrophages are well known for their ability to mediate antibody-dependent phagocytosis/immune signaling via engagement of Fc receptors expressed on them. We performed GSEA in macrophages from patients with SARS-CoV-2 infection, and compared macrophages between patients with mild and severe disease using KEGG gene sets. We acquired the top five significant gene sets resulting in upregulation of genes associated with TLR signaling on macrophages from patients with severe disease (Figure 3D). In addition, expression of activating Fc receptor genes (*FGCR1A, FCGR2A, FCGR3A, FCAR, FCMR*) were increased in macrophages from the severe group, compared with those from the mild group (Figure 3E). GSEA with gene sets for Fc receptor-mediated phagocytosis and signaling revealed significant upregulation of genes in macrophages from patients with severe disease (Figure 3F, S3). Taken together, these results indicated that the macrophages in the BAL fluids from patients with severe COVID-19 had more pro-inflammatory features and these might be induced by activation of Fc receptor-mediated signaling.

### Recruitment of Plasma Cells is Induced in Patients with Severe COVID-19

Plasma cells, differentiated from B cells, are professional antibody-producing cells during SARS-CoV-2 infection. We also analyzed plasma cells in the BAL fluids. The analysis revealed that the proportion and count of plasma cells in the patients with mild disease was slightly increased but was considerably more increased in the patients with severe disease (Figure 4A). In addition to accumulation of plasma cells, the GSEA result indicated that compared with patients with mild disease, plasma cells from patients with severe disease had up-regulated gene set, which is immunoglobulin complex (Figure 4B). These results suggested that plasma cells from patients with severe disease would have a relatively more vigorous response.

**Figure 4.**
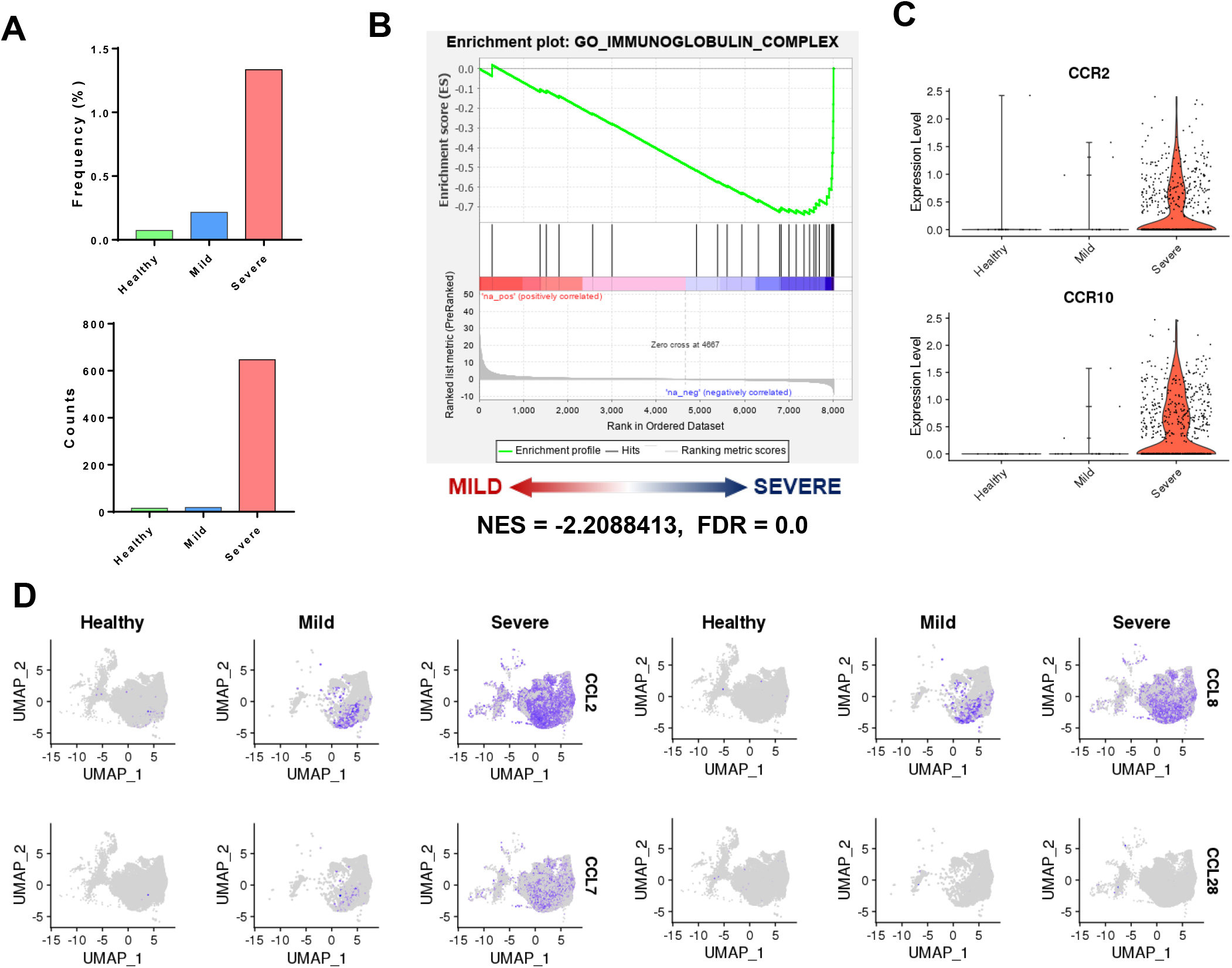
Accumulation of plasma cells in patients with severe COVID-19. (A) Proportions (up) and counts (down) of plasma cells among healthy controls and patients with mild or severe SARS-CoV-2 infection. (B) GSEA plot of DEGs in plasma cells from patients with mild vs severe disease, GO_IMMUNOGLOBULIN_COMPLEX gene set. (C) Expression of *CCR2* and *CCR10* on plasma cells from each sample. (D) UMAP plots of *CCL2, CCL7, CCL8,* and CCL28 gene expression on macrophages from healthy controls and from patients with mild and severe infection.

To examine factors associated with the accumulation of plasma cells in the BAL fluids of patients with severe disease, we analyzed the expression of chemokine receptor genes. Among chemokine receptors expressed in plasma cells (24,25), we found that *CCR2* and *CCR10* expression were dramatically increased in plasma cells of patients with severe disease (Figure 4C). Thus, we used feature plots to analyze the expression of chemokine genes, which are ligands for CCR2 and CCR10 (e.g., *CCL2, CCL7, CCL8* (for CCR2), and *CCL28* (for CCR10)), on macrophages (Figure 4D). These plots revealed that expression of *CCL2*, *CCL7*, and *CCL8*, but not *CCL28*, were considerably increased in the macrophages from the patients with severe disease. These results suggested that in patients with severe COVID-19 plasma cell accumulation was induced by increased CCR2-associated expression of chemokines on macrophages.

## Discussion

In this study, focusing on B cells, we analyzed different characteristics of cells from BAL fluids of patients with mild or severe symptoms and clinical signs. We based the analysis on single-cell RNA sequencing data. We found that the B cells from patients with mild disease had different characteristics compared to those from patients with severe disease. These results of GSEA and chemokine receptor gene expression indicated that B cells from the mild group were much closer to naïve/memory B cells, compared with those from the severe group. This suggested that B cells from the severe group had features more similar to plasma cells. Likewise, a transcriptomic analysis of peripheral blood mononuclear cells (PBMCs) from COVID-19 patients previously showed that the proportion of naïve/memory B cells in late time point of recovery stage was higher than in early time point (26). In addition, B cells from the severe group also had higher expression of Ig genes and activation characteristics. Therefore, our analysis suggested that B cell activation might occur larger during severe disease.

The up-regulation of genes related to TNF-α signaling pathway occurred in B cells from severe group samples, compared with those from mild group samples. This result suggested that BAFF and APRIL, members of TNF superfamily, might affect B cells with SARS-CoV-2 infection as well. Macrophages from patients with severe disease had higher expression of *TNFSF13B (BAFF)* but much lower expression of *TNFSF13 (APRIL)*, compared with those from patients with mild disease. In a similar vein, a previous study showed that, in protein level, elevated BAFF and reduced APRIL in serum from patients with active SARS-CoV-2 infection compared to recovered patient (12). Both BAFF and APRIL promote B cell survival, differentiation into plasma cells, and antibody production. However, if these two molecules produce heterotrimers, they have decreased potency to induce B cell proliferation and compete with BAFF homotrimer. This finding suggests that APRIL can negatively regulate BAFF activity (27). Moreover, APRIL promotes anti-inflammatory cytokine interleukin-10 production and regulatory functions in human B cells (28). Therefore, lower expression of APRIL in macrophages from severely affected patients might induce B cell activation and differentiation.

As found in a previous study (5), we found that macrophages from patients with severe disease had more features associated with the inflammatory response, compared with patients with mild symptoms. In this analysis, the GSEA results for comparison of pathways between macrophages from patients with mild or severe disease indicated that macrophages from severely affected patients up-regulated genes related to the TLR signaling pathway. Activating Fc receptor gene expression was also greater on macrophages from the severe group. Additional GSEA results indicated that compared with samples from the mild group, up-regulation of genes related to Fc receptor signaling occurred in macrophages from samples from the severe group. Furthermore, our analysis showed the increased expression of *CCL2* on macrophages from severe patients compared to mild patients, suggesting the possibility of ADE occurrence. Also, the accumulation of plasma cells was induced in patients with severe symptoms, and those cells were highly expressed *CCR2* which is the receptor for CCL2.

The previous study observed the ADE phenomenon in the SARS-CoV-1 infection, showing that neutralizing antibody activities were higher in deceased patients than recovered patients and the production of antibodies in deceased patients were elicited faster than in recovered patients (29). This phenomenon is known to exacerbates viral infections by antibody-dependent mechanisms such as phagocytosis with Fc receptors and TLR signaling (30). Also, several studies have shown the association between higher titers of antibodies and severe symptoms in SARS-CoV-2 infection (1,13). Our analysis showed increased expression of Ig genes and activation of B cells, accumulation of plasma cells, and up-regulation of genes related to Fc receptor signaling in macrophages from patients with severe COVID-19, which are associated with antibody-dependent enhancement phenomenon. Our reanalysis of transcriptomic data suggested a possibility of ADE phenomenon occurrence in macrophages from patients with severe disease, leading to hyper-inflammation.

In conclusion, our study showed different characteristics of B cells from BAL fluids between patients with mild and severe COVID-19 and the accumulation of CCR2-expressed plasma cells in severe group. Also, we figure out different expressions of BAFF and APRIL genes on macrophage, which involve in the regulation of B cells, between mild and severe group. However, our study is based on a small cohort of patients with individual variability, so the result may be biased. Therefore, larger amounts of samples may allow to better characterize B cells and relevant regulations.

## Acknowledgments

The authors thank Dr. Zheng Zhang, who provided valuable raw RNA sequencing data of patients with COVID-19. This work was supported by the National Research Foundation of Korea (NRF-2018M3A9H3024611, NRF-2019M3A9A8067236), and Mobile Clinic Module Project funded by KAIST, Republic of Korea.

## Author Contributions

C.W.K. and H.K.L designed and performed the study. C.W.K. J.E.O. and H.K.L. conceived the study, analyzed the data, and wrote the manuscript.

