## Supplementary material for "Single cell transcriptomic re-analysis of immune cells in bronchoalveolar lavage fluids reveals the correlation of B cell characteristics and disease severity of patients with SARS-CoV-2 infection": Supplemetary figures and table

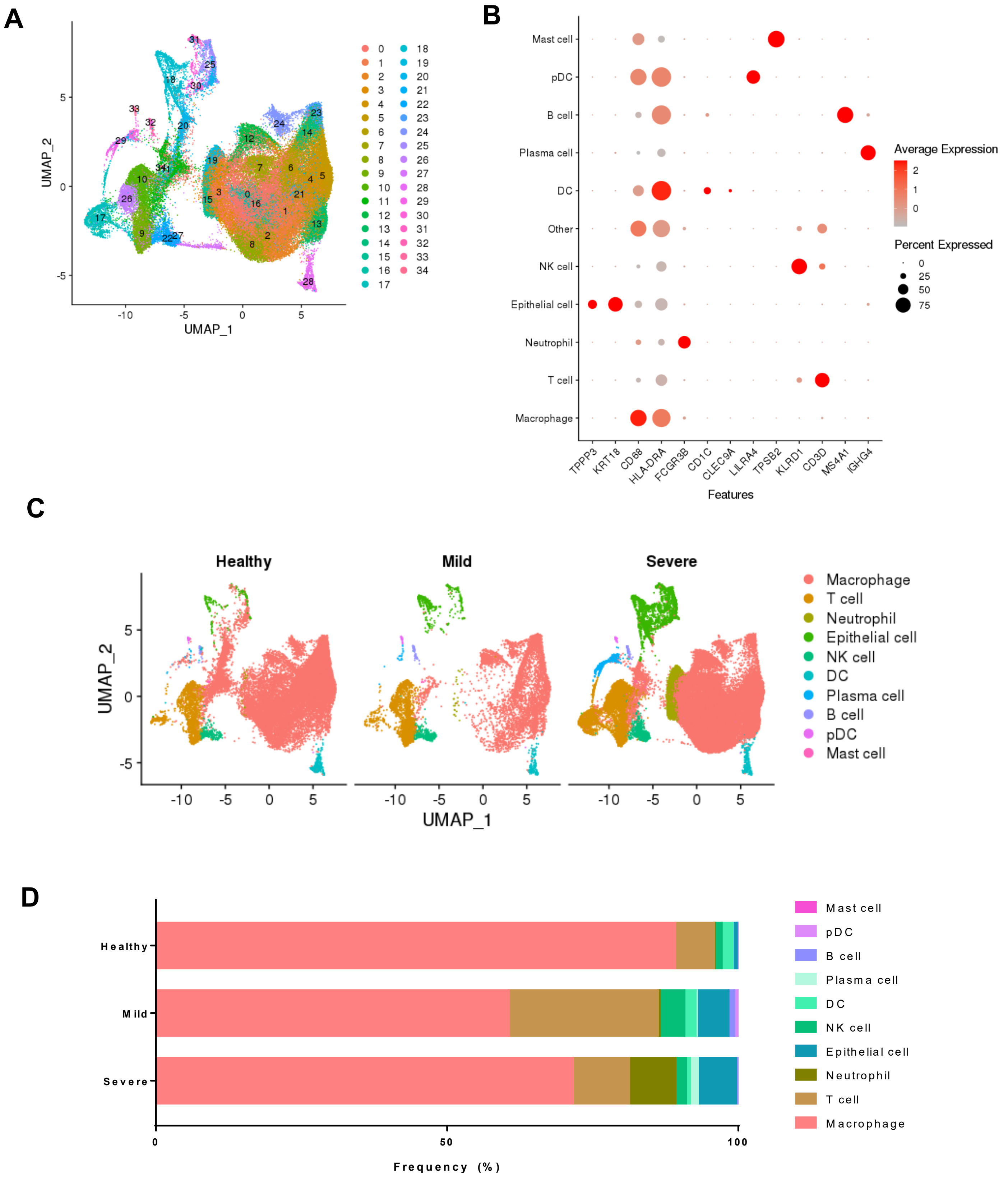


**Figure S1. The composition of immune cells in bronchoalveolar lavage (BAL) fluids in COVID-19 patients**

(A) UMAP plot of immune cell types in BAL fluids in patients with COVID-19 infection. (B) Dot plot showing expression of marker genes by different cell clusters. (C) UMAP plots showing cell clusters for healthy controls, and patients with mild and severe infection. (D) The proportions of cell clusters for each group.

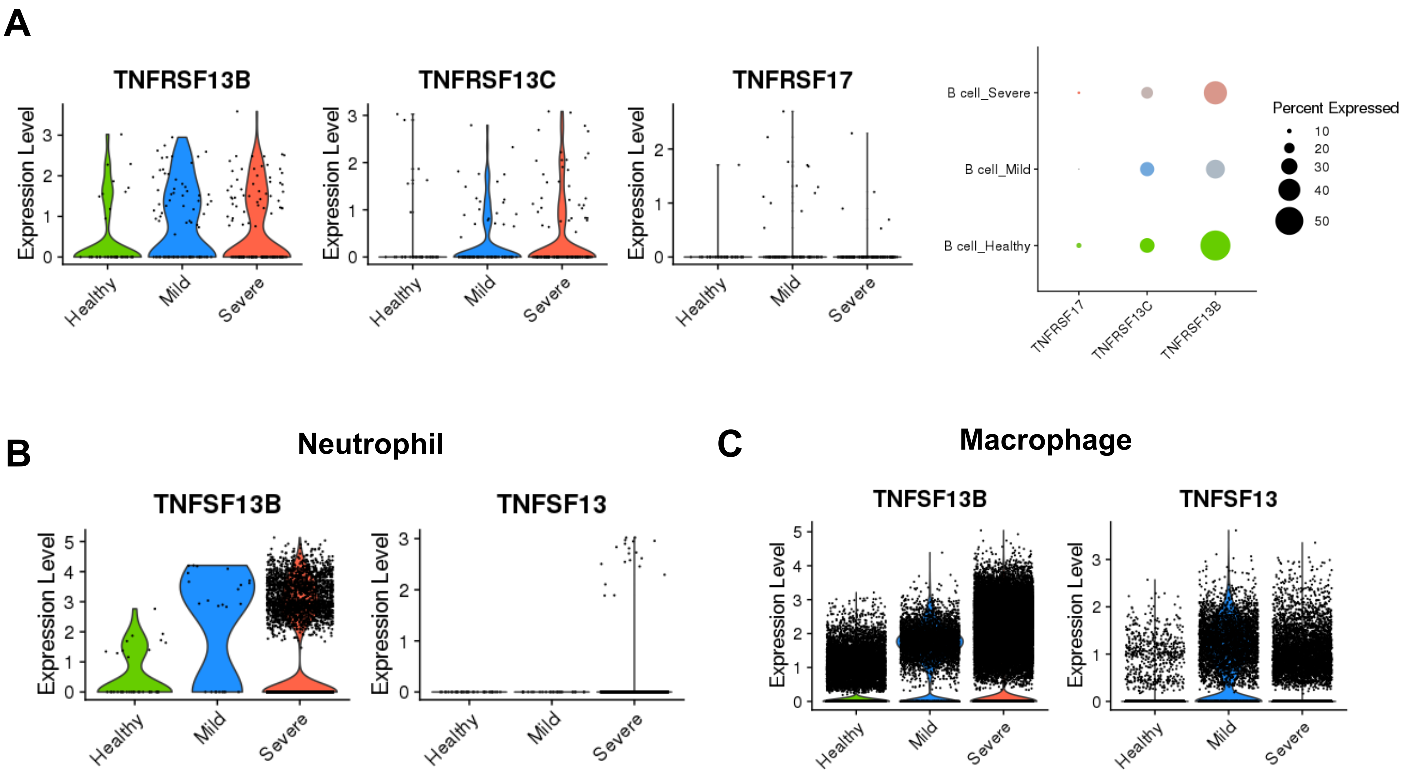


**Figure S2. The expression of BAFF, APRIL and their receptor genes**

(A) Violin plots and dot plot showing expression of receptor genes (*TNFRSF13B, TNFRSF13C, TNFRSF17*) for BAFF and APRIL from each group. (B, C) Expression of *TNFSF13B* and *TNFSF13* on (A) neutrophils and (B) macrophages from each group. (C)

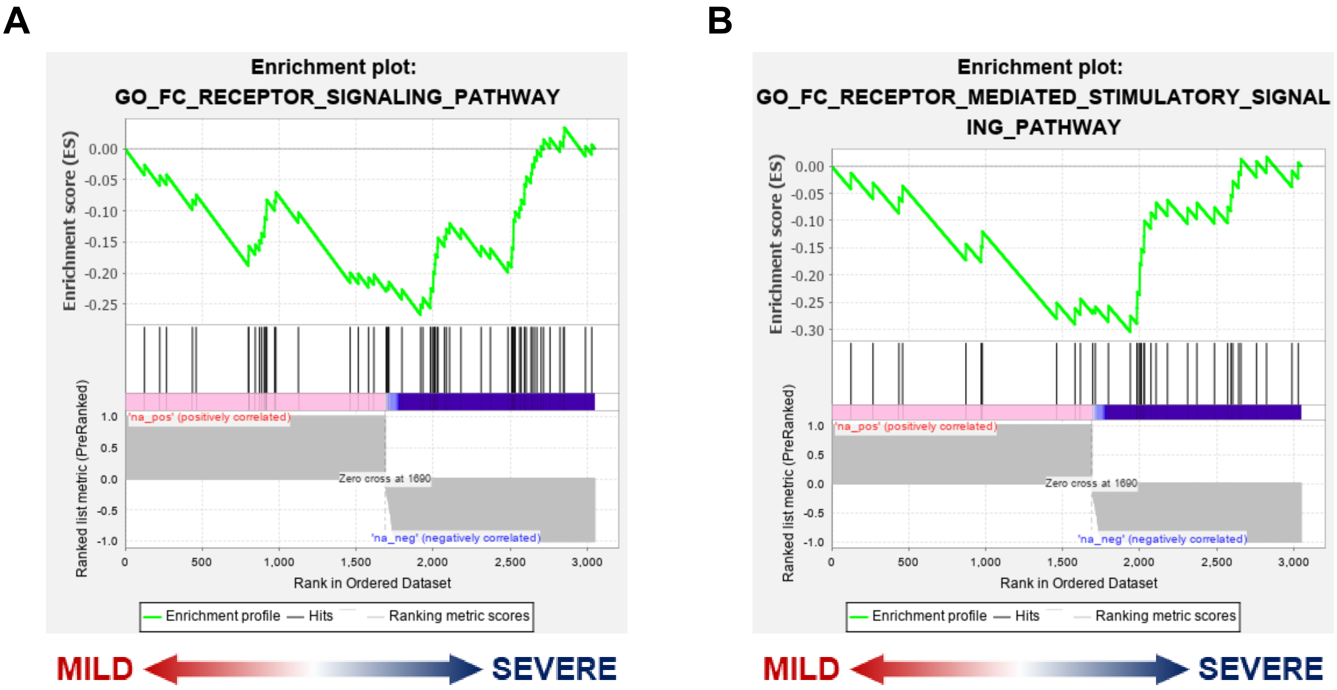


**NES = -2.4703496, FDR = 0.0**

**NES = -2.1226537, FDR = 0.006810443**

**Figure S3. Fc receptor-associated pathways in macrophages of severe patients**

(A, B) GSEA plots of DEGs in macrophages of mild vs severe patients with (A) the GO_FC _RECEPTOR_SIGNALING_PATHWAY gene set and (B) the GO_FC_RECEPTOR_MEDIATED _STIMULATORY _SIGNALING_PATHWAY gene set.

**Supplementary Table 1. Clinical data of COVID-19 patients examined by scRNA-seq**

| **Group** | **Mild** | | | **Severe** | | | | | |
|---|---|---|---|---|---|---|---|---|---|
| **Patient** | 1 | 2 | 3 | 1 | 2 | 3 | 4 | 5 | 6 |
| **Age / Gender** | 36 / M | 37 / F | 35 / M | 62 / M | 66 / M | 63 / M | 65 / F | 57 / F | 46 / M |
| **Severity** | Moderate | Moderate | Moderate | Severe | Critical | Critical | Critical | Critical | Critical |
| **Time to sampling post symptom onset (day)** | 11 | 9 | 13 | 11 | 18 | 14 | 25 | 8 | 12 |
| **Outcome** | Cured | Cured | Cured | Cured | Death | Death | Cured | Cured | Cured |
| **Chronic basic disease** | None | None | None | None | Hypertension | Sleep apnea | Diabetes | None | None |
| **Medication history** | None | None | None | None | Amlodipine Besylate | None | None | None | None |
